# Humanization and Expression of IgG and IgM Antibodies in Plants as Potential Diagnostic Reagents for Valley Fever

**DOI:** 10.1101/2022.04.27.489777

**Authors:** Collin Jugler, Francisca J. Grill, Thomas E. Grys, Douglas F. Lake, Qiang Chen

## Abstract

Monoclonal antibodies (mAbs) are important proteins used in many life science applications, from diagnostics to therapeutics. High demand for mAbs for different applications urges the development of rapid and reliable recombinant production platforms. Plants provide a quick and inexpensive system for producing recombinant mAbs. Moreover, when paired with an established platform for mAb discovery, plants can easily be tailored to produce mAbs of different isotypes against the same target. Here, we demonstrate that a hybridoma-generated mouse mAb against chitinase 1 (CTS1), an antigen from *Coccidioides* spp., can be biologically engineered for use with serologic diagnostic test kits for coccidioidomycosis (Valley Fever) using plant expression. The original mouse IgG was modified and recombinantly produced in plants as IgG and IgM isotypes with human kappa, gamma, and mu constant regions. The two mAb isotypes produced in plants were shown to maintain target antigen recognition to CTS1 using similar reagents as the Food and Drug Administration (FDA)-approved Valley Fever diagnostic kits. As none of the currently approved kits provide antibody dilution controls, humanization of antibodies that bind to CTS1, a major component of the diagnostic antigen preparation, may provide a solution to the lack of consistently reactive antibody controls for Valley Fever diagnosis. Furthermore, our work provides a foundation for reproducible and consistent production of recombinant mAbs engineered to have a specific isotype for use in diagnostic assays.

## Introduction

Antibodies are a leading class of biological molecules utilized in numerous research, diagnostic, and treatment settings. Monoclonal antibodies (mAbs) have a strong research and development interest for therapeutic purposes due to a highly profitable market, predicted to be upwards of $300 billion by 2025^1^. The market for antibodies as research reagents or diagnostic tools is estimated to reach levels of $6.3 billion by 2027 and $42.56 billion by 2029, respectively^2,3^. Many antibodies used for these latter purposes are polyclonal in nature, allowing for more economical production. However, polyclonal antibodies may introduce a large degree of batch-to-batch variation, causing reproducibility issues^4,5^. These issues can be overcome by identifying sequences of mAbs for targets of interest, thereby allowing recombinant production processes to replace mammalian serum collection as an antibody source. Strong market projections, in parallel with the reproducibility crisis, urges innovation in both identifying new mAbs, as well as improving production methods for sustainable, reliable sources for reagents.

Generating mAbs to targets of interest can be accomplished in several ways. Isolation of human B cells, followed by antigen-specific sorting and single-cell variable region sequencing to identify high affinity and specific antibodies is a high throughput method employed when convalescent individuals are available^6^. However, this method may not be useful for discovering antibodies against antigens that individuals may not been exposed to and thus, do not have B cells that have undergone proper affinity maturation. Also, this method is not cost-effective, requiring cutting-edge technology for successful mAb generation, limiting its accessibility. Phage display is another technology that has generated useful mAbs for therapeutic purposes, although it also has limitations, including libraries not representing all antibodies and potential mispairing of variable heavy and light regions, compared to *in vivo* generation of antibodies^7,8^. The traditional and widely used method of hybridoma generation following mouse immunization can generate highly pure, full length mAbs that can be continuously cultured for production^9^. However, even hybridoma technology can suffer from instability of the fused cells and there is constant risk of contamination, which can limit sustainable production of a consistent mAb. Furthermore, traditional hybridoma-derived mAbs are of mouse origin, making them unsuitable for human applications. This limitation also pertains to certain *in vitro* diagnostic applications. For example, hybridoma-produced murine mAbs cannot be used as positive controls in commercial assays for diagnosis of antibody responses to pathogens in human patients, as the secondary antibodies (usually anti-human IgG or IgM) cannot recognize the control mAbs of mouse origin.

Recombinant protein expression in plants is an alternative to other expression systems that has gained momentum in recent decades. MAbs against a diverse array of targets have been generated and characterized, with some being produced under current Good Manufacturing Practices (cGMP) as proof-of-principle that plants can be a viable alternative to mammalian-made biopharmaceuticals^10,11^. There are several advantages of using plants as mAb production hosts. Plants do not require sterile growth conditions or expensive growth media, making them a more cost-effective production method compared to Chinese hamster ovary (CHO) or hybridoma-based methods, while also greatly reducing the risk of contamination of mammalian viruses or other pathogens^12,13^. Once the variable gene sequences of a mAb of interest are defined, simple cloning procedures followed by agroinfiltration^14,15^ allows the rapid generation of high levels of mAbs in plants. The flexibility of plant-based expression systems has allowed the production of multiple mAbs and mAb variants for a diversity of applications, including engineering the same Fab region to different antibody isotypes and subtypes, and mAbs for diagnostic purposes^16–21^.

Serological diagnostic tests are used to help identify many different diseases in patients. The specificity of antibodies generated from the immune response against pathogens can be detected by enzyme immunoassays (EIA) to inform the infection status. Coccidioidomycosis (Valley Fever) is a fungal respiratory disease caused by *Coccidioides* spp^22^. The primary clinical diagnostic test for Valley Fever detects the antibody response in patient serum to the fungi^23^. Specifically, patient IgG and IgM against Valley Fever are detected in an EIA by reactivity to *Coccidioides* spp. antigen coated in individual wells on a plate^24^. Clinical laboratories performing serology tests using Food and Drug Administration (FDA)-approved kits for diagnosis of Valley Fever are required to run dilution controls when testing patient sera, per Clinical Laboratory Improvement Amendments (CLIA) regulations^25^, ensuring that patient samples and controls are diluted in the same manner while performing the assay. Since human IgG and IgM dilution controls are not provided in the diagnostic serology kits, human sera that demonstrated previous IgG and/or IgM reactivity to *Coccidioides* spp. antigen must be identified, stored and re-used with the reagents in the diagnostic kit every time the test is run to comply with federal requirements. The supply of positive human sera is uncertain, and the inherent variability of human sera may lead to inconsistent diagnostic results. The lack of a consistent control presents a major challenge to all clinical laboratories that perform serologic testing for Valley Fever and provides an opportunity to generate human IgG and IgM reagents that can be used as dilution controls with the serologic test kits.

Here, we present the development of a *Coccidioides* spp. antigen-specific mAb produced by murine hybridomas and its conversion to humanized chimeric IgG and IgM isotypes by using a plant-based expression system. The two mAb human isotypes are produced efficiently in plants. Furthermore, they effectively recognize their target, chitinase 1 (CTS1), from *Coccidioides* spp. using similar reagents to those used clinically in Valley Fever diagnostic kits. This study may provide a solution to a particular “pain-point” in clinical laboratories that run serologic tests for the diagnosis of coccidioidomycosis. This may also set a foundation for using plants to provide various human antibody isotypes as human diagnostic reagents, inexpensively and quickly with consistent quality.

## Methods & Materials

### Antigen preparation and mouse immunization

Antigen was prepared as a culture filtrate of the mycelial phase of *Coccidioides* spp. as previously detailed^26^. Mycelial culture filtrate (MCF) was concentrated using an Amicon centrifugal filter unit (Millipore) and the total protein concentration was measured by bicinchoninic acid (Thermo Scientific). MCF was mixed with Magic Mouse adjuvant (Creative Diagnostics) at 1:1 volume and used to immunize one BALB/c mouse under an Institutional Animal Care and Use Committee–approved protocol at Mayo Clinic (50 µg per dose). Subcutaneous (SQ) immunization was repeated with half the dose of MCF at 3-week intervals. Antibody titers to MCF were monitored by indirect ELISA and when an acceptable level was reached (>1:32,000), a final SQ boost was given, and the mouse was sacrificed three days later for hybridoma generation.

### Generation and purification of mAbs

Hybridomas were generated using a standard technique^9^. Briefly, the spleen was excised and processed into a single-cell suspension, followed by lysis of erythrocytes with 1X red blood cell lysis buffer (Invitrogen). The remaining 4 × 10^7^ splenocytes were fused with 8 × 10^7^ murine myeloma cells (P3X63Ag8.653) using 1 mL of 50% polyethylene glycol solution (Sigma-Aldrich). Fused cells were resuspended in 20% fetal bovine serum complete Dulbecco’s modified Eagle medium with hypoxanthine-aminopterin-thymidine selection supplement (Sigma-Aldrich) and plated in 96-well plates at 5 × 10^4^ splenocytes per well. Plates were incubated for ten days at 37°C in a 5% CO_2_ incubator. Culture supernatant was tested for antibodies to MCF by indirect ELISA. Cells positive by ELISA were subcloned by limiting dilution and screened using the same method after ten days. Positive subclones were expanded and supernatant was purified by protein A/G chromatography (Thermo Scientific). Purified mAbs were evaluated for specificity by Western blotting. One mAb, 4H2, bound to a known antigen in MCF, CTS1, and was pursued for gene rescue.

### ELISA for Hybridoma Screening

Purified MCF was coated on an ELISA plate at 20 µg/mL. Then, the plate was washed with 1X phosphate-buffered saline + 0.05% Tween-20 (PBST) three times and blocked in 1% bovine serum albumin in phosphate-buffered saline (BSA-PBS). For monitoring mouse antibody titers, mouse serum was diluted in 1% BSA-PBS to various dilutions and allowed to incubate on the ELISA plate for 1 hour. For testing hybridoma cultures, neat supernatant was added directly to the blocked ELISA plate and incubated for 1 hour. After primary antibody incubation, plates were washed three times with PBST then incubated with horseradish peroxidase (HRP)-conjugated goat-anti mouse IgG antibody (1:5000 dilution, Jackson ImmunoResearch, Cat. No. 115-035-071) for 1 hour. Plates were washed four times with PBST and TMB substrate was added. After color development, the reaction was stopped with 0.16M sulfuric acid and absorbance was read at 450nm.

### Mouse immunoglobulin gene rescue

Cell lysate of mAb 4H2 was prepared using the Cells-to-cDNA™ II Kit (Invitrogen). The cell lysate was subsequently used to synthesize cDNA using a reverse transcription reaction with reagents included in the kit. The resultant cDNA was subject to polymerase chain reaction (PCR) amplification of unknown mouse immunoglobulin light chain (LC) and heavy chain (HC) variable genes using previously published degenerate primers^27^. Each PCR reaction contained 2 µL cDNA, 0.5 µM 5’ and 3’ primers, 25 µL Phusion Flash High-Fidelity PCR Master Mix (Thermo Scientific), and ultra-pure water to bring the reaction volume to 50 µL. Cycling conditions were as follows: initial denaturation at 98°C for 10 seconds followed by 30 cycles of a three-step program (98°C, 1 second; 55°C, 5 seconds; 72°C, 6 seconds), and a final extension at 72°C for 1 minute. Amplification was confirmed by running 8 µL of the PCR reaction on a 1% agarose gel. The remaining 42 µL of the PCR product were subject to DNA cleanup using the QIAquick® PCR Purification Kit (Qiagen) and the approximate concentration of DNA was determined using a NanoDrop spectrophotometer (Thermo Scientific). Cleaned DNA from each PCR reaction was cloned into pcDNA™3.1V5/HisA vector (Thermo Scientific) using the Cold Fusion Cloning Kit (System Biosciences). DNA from resultant clones was extracted using the QIAprep®Spin Miniprep Kit (Qiagen) and sent for Sanger sequencing.

### Human IgG and IgM Chimera Construction

Primers were designed to add a plant-specific secretion signal peptide to murine variable regions of the LC and HC clones. The resulting PCR products were then digested and ligated onto a human kappa chain backbone, along with a human IgG_1_^28^ and a human mu chain backbone^17^, respectively, before being cloned into a plant expression vector as described^29^. Positive clones containing the human chimeric IgG and IgM constructs were transformed in *Agrobacterium tumefaciens* and verified by PCR.

### Agroinfiltration and Temporal Expression in *Nicotiana benthamiana*

Transgenes were introduced into *N. benthamiana* leaves by agroinfiltration as previously described^14,30^ and leaves expressing the recombinant IgG or IgM were collected for temporal expression analysis. Leaves collected in the range of 2 – 9 days after transgene introduction were homogenized in extraction buffer (1X PBS containing 1mM ethylenediaminetetraacetic acid [EDTA], 2mM phenylmethylsulfonyl fluoride [PMSF], and 10 mg/mL sodium L-ascorbate) at a 1:1.5 ratio of fresh leaf weight (FLW) to buffer volume. An ELISA plate that had been coated overnight at 4°C with a goat anti-human kappa chain antibody (Southern Biotech, Cat. No. 2060-01) at 2 µg/mL, was washed four times with PBST, followed by blocking with 5% dry milk dissolved in PBST (DM-PBST). Dilutions in DM-PBST of homogenized leaf samples and an isotype control of known concentration were added to the plate and incubated for 1 hour at 37°C. After washing four times, a goat anti-human IgG-HRP antibody (1:4000 dilution, Southern Biotech, Cat. No. 2040-05) or a goat anti-human IgM mu chain-HRP (1:5000 dilution, Abcam, Cat. No. ab97205) was diluted in 5% DM-PBST and added to the plate for 1 hour at 37°C. After an additional four washes, TMB substrate was added to the plate and a 1M sulfuric acid stop solution was used prior to reading absorbance at 450nm.

### Plant-Made mAb Purification and SDS-PAGE Analysis

*N. benthamiana* leaves expressing either the humanized 4H2 IgG or IgM were harvested and homogenized in the same extraction buffer described above for temporal analysis. The protein extract containing IgG or IgM was processed as described^29,31^, followed by Protein A (Cytiva) or human anti-mu chain (Sigma-Aldrich, Cat. No. A9935) affinity chromatography. Eluted mAbs were further concentrated and buffer exchanged into 1X PBS, pH 7.4 by ultrafiltration in an Amicon centrifugal filter unit (Millipore). Purified mAbs were then subjected to SDS-PAGE and Coomassie Blue R-250 staining under both reducing and non-reducing conditions on a Mini-Protean TGX 4-20% gel (Bio-Rad).

### Western Blotting

Western blots were performed as described previously^29^. In brief, to validate the plant-made mAbs identity, plant-made 4H2 IgG (P-4H2 IgG) or plant-made 4H2 IgM (P-4H2 IgM), were subjected to SDS-PAGE under reducing and non-reducing conditions and transferred onto polyvinylidene difluoride (PVDF, Bio-Rad) membranes. Membranes were blocked with 5% DM-PBST for 1 hour, followed by washing with PBST. Goat anti-human kappa-HRP (1:5000 dilution, Southern Biotech, Cat. No. 2060-05), goat anti-human IgG-HRP (1:10,000, Southern Biotech, Cat. No. 2040-05) or goat anti-human IgM mu chain-HRP (1:5000 dilution, Abcam, Cat. No. ab97205) were diluted in 5% DM-PBST and incubated for 1 hour. After washing with PBST, membranes were developed using with SuperSignal® West Pico Chemiluminescent Substrate (Thermo Scientific) according to manufacturer’s instructions and images were taken with an ImageQuant instrument.

For validation of the plant-made mAbs’ target recognition, recombinant CTS1 and *Coccidioides* MCF were separated under reducing conditions on 12% polyacrylamide gels and transferred to PVDF membrane. Following blocking in 1% BSA-PBST, either the parental murine 4H2 IgG (M-4H2 IgG), P-4H2 IgG, or P-4H2 IgM were diluted to 500 ng/mL in 1% BSA-PBS and incubated for 1 hour. The membranes were then probed with the appropriate secondary antibody [1:10,000 dilution, goat-anti-mouse IgG Fc-specific-HRP, Jackson ImmunoResearch, Cat. No. 115-035-071; 1:10,000 dilution, goat-anti-human IgG Fc-specific-HRP, Sigma-Aldrich, Cat. No. A0170; 1:10,000 dilution, goat-anti-human IgM Mu-HRP, Caltag Cat. No. H15007], followed by a 1-hour incubation. After washing with PBST, a KPL TrueBlue Peroxidase substrate (SeraCare) was added, followed by image capture.

### Antigen Binding ELISA by Chimeric IgG and IgM

To evaluate P-4H2 IgG and P-4H2 IgM antigen binding, recombinant CTS1 was coated on an ELISA plate at 2 µg/mL overnight at 4°C. Then, the plate was washed with 1X PBST three times and blocked in 1% BSA-PBS. MAb isotypes were tested at 1 µg/mL and diluted two-fold in 1% BSA-PBS. After 1 hour of incubation, plates were washed three times with PBST then incubated with HRP-conjugated anti-human secondary antibody [1:20,000 dilution, goat-anti-human IgG Fc-specific-HRP, Sigma-Aldrich, Cat. No. A0170; 1:5,000 dilution, goat-anti-human IgM Mu-HRP, Caltag Cat. No. H15007]. Hybridoma-produced parent mAb was used as a positive control, with an HRP-conjugated anti-mouse secondary antibody [1:20,000 dilution, goat-anti-mouse IgG Fc-specific-HRP, Jackson ImmunoResearch, Cat. No. 115-035-071]. Plates were washed four times with PBST and TMB substrate was added. The reaction was stopped with 0.16M sulfuric acid and absorbance was read at 450nm.

## Results

### Engineering and Expression of Coccidioides-Specific Human of IgG and IgM Chimeras in Nicotiana benthamiana

A mAb of interest, 4H2, was identified by immunizing mice with MCF from *Coccidioides* spp., followed by hybridoma generation and gene rescue. The resulting murine variable heavy region was genetically fused onto both human IgG_1_ and IgM heavy constant chains, in parallel with the variable light region being grafted onto a human kappa constant chain. Gene constructs for both humanized IgG and IgM were introduced into leaves of *N. benthamiana* and recombinant expression of the 4H2 IgG and IgM chimeras was monitored over time by an anti-human IgG or IgM ELISA. Peak expression of P-4H2 IgG was observed at day 7 after agroinfiltration, reaching 39.95 ± 2.5 µg/g FLW (**Figure 1A**). P-4H2 IgM reached peak expression of 33.3 ± 1.19 µg/g FLW at day 3 after introduction of the transgenes (**Figure 1B**).

**Figure 1:**
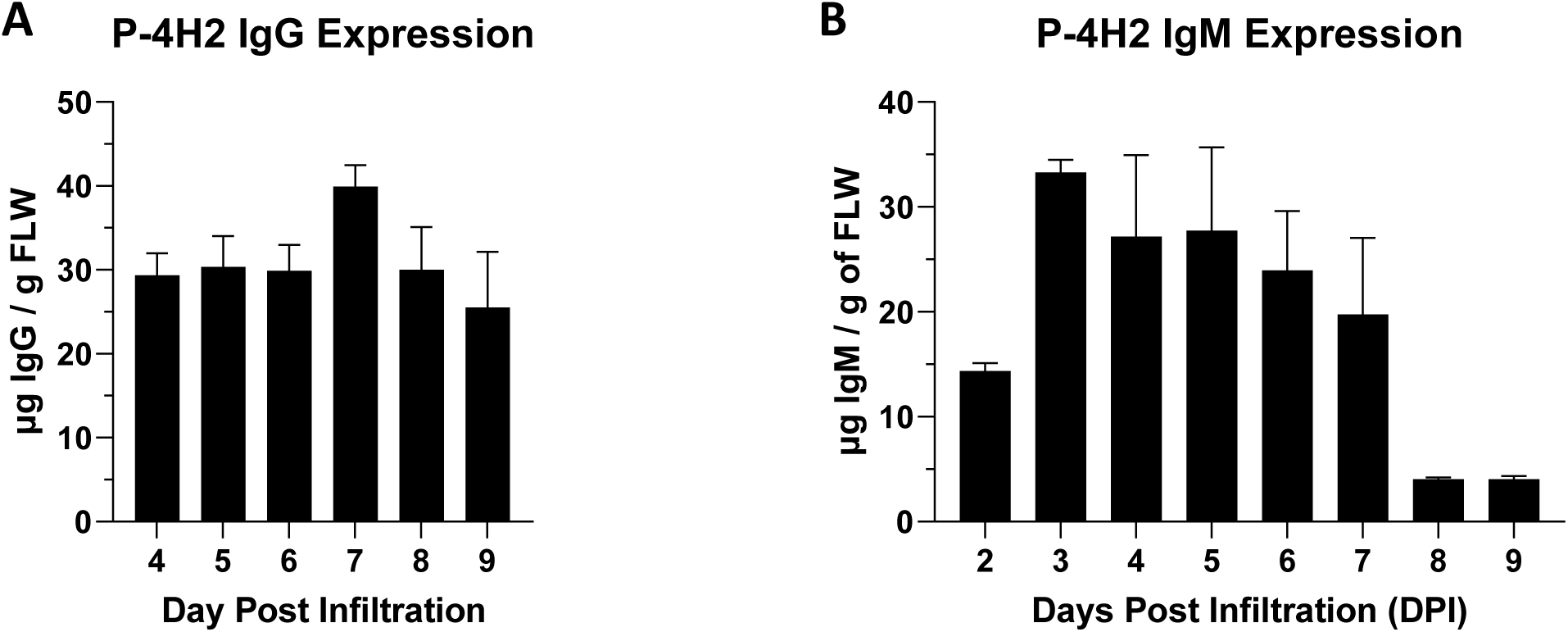
Expression of 4H2 Isotypes in *Nicotiana benthamiana* leaves. Temporal expression of P-4H2 IgG (**A**) and P-4H2 IgM (**B**) human chimeras was analyzed by sandwich ELISA that detects IgG or IgM containing both human kappa light chain and either human gamma or human mu heavy chain from total soluble plant protein extracts. Mean ± SEM from two independent experiments is plotted.

### Recombinant Chimeric IgG and IgM Purification and Characterization

P-4H2 IgG was purified to a high degree of homogeneity by Protein A affinity chromatography, comparable to a control IgG mAb (**Figure 2A**). The recombinant P-4H2 IgM was purified by affinity chromatography with a goat anti-human mu chain agarose resin (**Figure 2B**) and more bands were observed than the IgG isotype. To verify the identity of the observed bands from affinity purified P-4H2 IgG and P-4H2 IgM, western blotting was performed. The bands observed with SDS-PAGE analysis of the IgG isotype indeed correspond to both human kappa chain at ∼25 kDa (**Figure 3A and 3C**), as well as human gamma chain at ∼50 kDa (**Figure 3B**). Likewise, the bands observed for the purified P-4H2 IgM represent the same human kappa chain (**Figure 4A and 4C**), as well as the human mu chain (**Figure 4B**), with an expected size of ∼60-65 kDa. Western blot analysis also revealed that the major protein bands observed in the purified, P-4H2 IgM falling between 50 and 37 kDa represent either proteolysis of intact human mu chain or a truncated mu chain, in addition to the expected band at ∼60-65 kDa (**Figure 2B and 4B**). As expected, both P-4H2 IgG and P-4H2 IgM under non-reducing conditions are composed of two heavy and two light chains, for monomeric sizes of ∼150 kDa (**Figure 3C**) and ∼170 kDa (**Figure 4C**), respectively.

**Figure 2:**
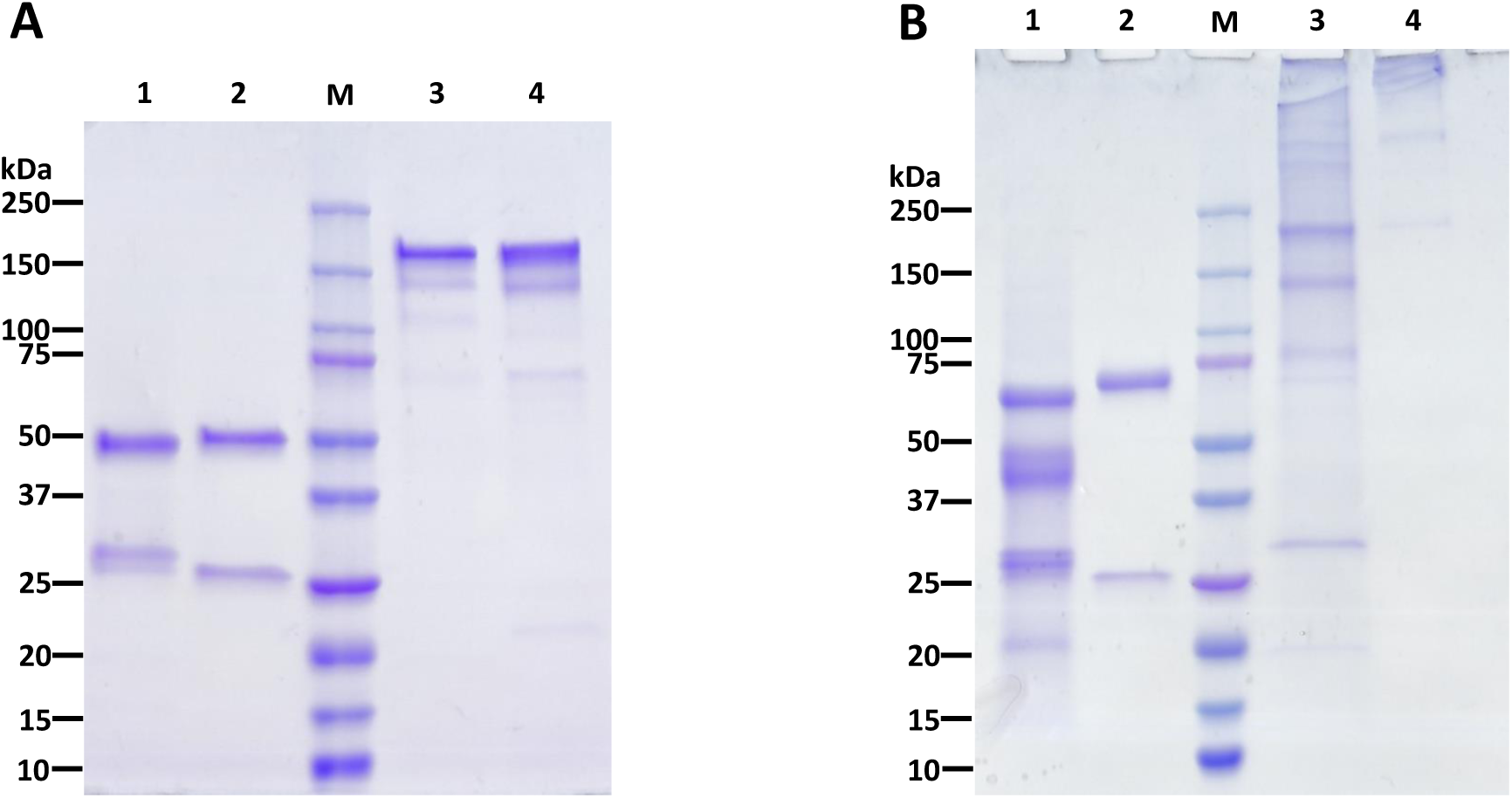
Purification of P-4H2 IgG and P-4GH2 IgM from *N. benthamiana* leaves. Affinity-purified, humanized P-4H2 IgG (**A**) and P-4H2 IgM (**B**) were separated by SDS-PAGE on 4-20% polyacrylamide gels under reducing and non-reducing conditions. One representative of multiple experiments is shown. Lanes 1 and 3, plant-made mAb isotypes. Lanes 2 and 4, control human mAb isotypes. M, molecular weight marker.

**Figure 3:**
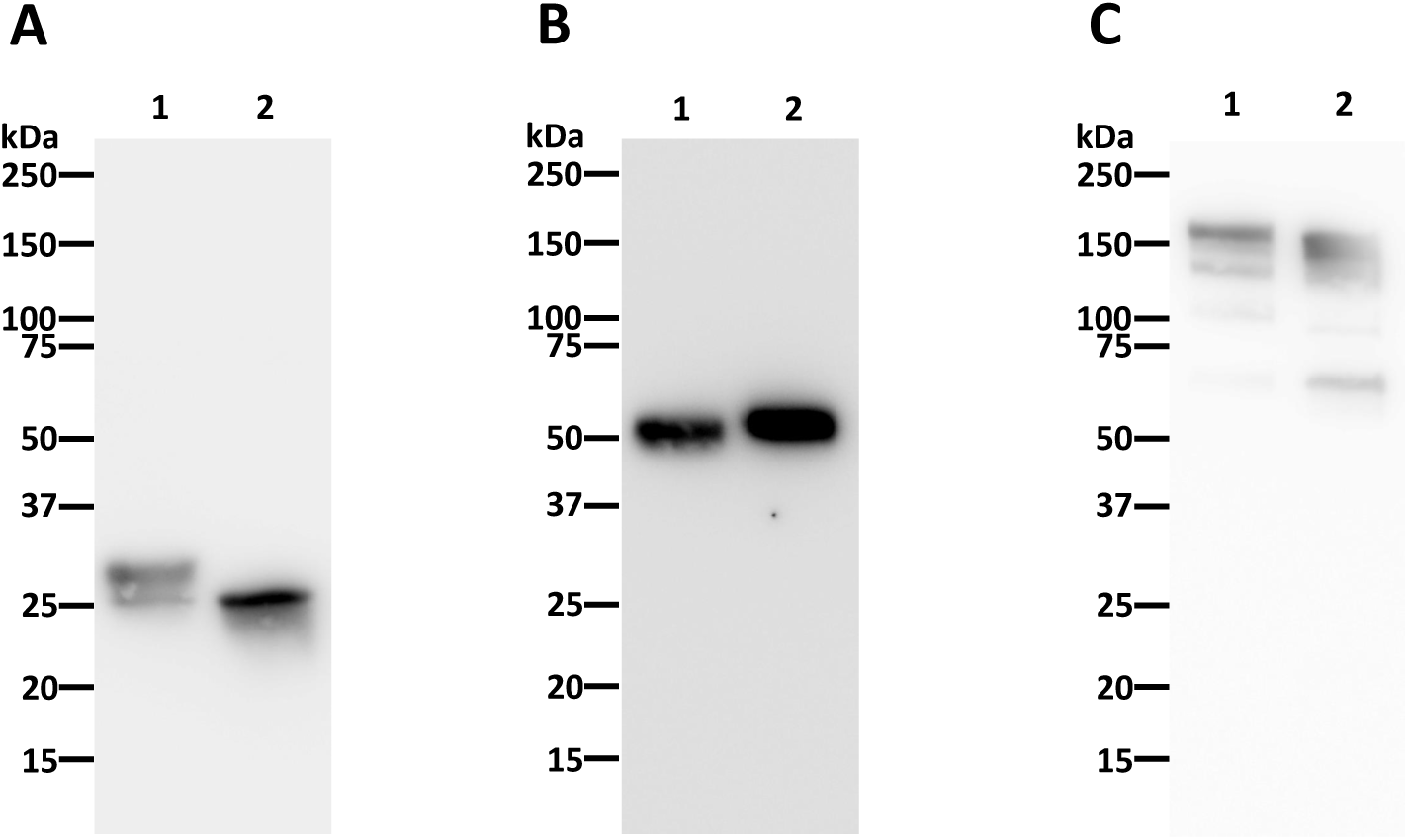
Assembly of 4H2 IgG purified from *N. benthamiana* leaves. Purified recombinant P-4H2 IgG was separated under reducing (**A** and **B**) and non-reducing (**C**) conditions by SDS-PAGE and proteins was transferred to PVDF membranes. Proteins were then detected with antibodies specific for human kappa chain (**A** and **C**) or human gamma chain (**B**) to verify protein identity and assess IgG assembly. Lane 1, Humanized, P-4H2 IgG, Lane 2, Human kappa IgG control.

**Figure 4:**
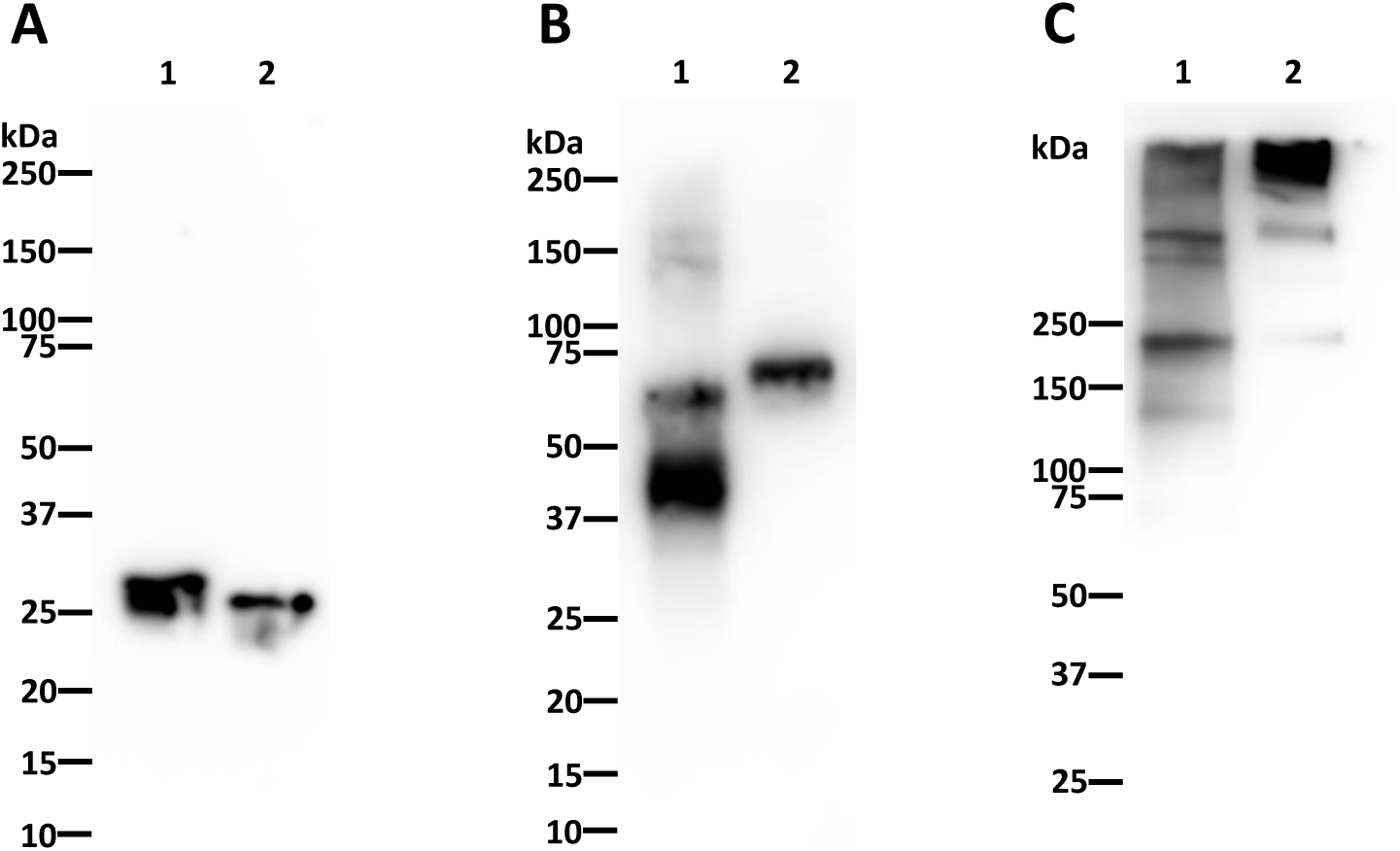
Assembly of 4H2 IgM purified from *N. benthamiana* leaves. Purified recombinant P-4H2 IgM was separated under reducing (**A** and **B**) and non-reducing (**C**) conditions by SDS-PAGE and proteins was transferred to PVDF membranes. Proteins were then detected with antibodies against human kappa chain (**A** and **C**) or human mu chain (**B**) to verify protein identity and assess IgM assembly. Lane 1, Humanized, P-4H2 IgM, Lane 2, Human kappa IgM control.

To ensure the human IgG and IgM chimeras produced in plants maintained antigen binding specificity of M-4H2 IgG, P-4H2 IgG and P-4H2 IgM were also tested for binding to recombinant CTS1 and MCF. As determined by indirect ELISA, both recombinant isotypes of the chimeric mAb bound to recombinant CTS1 in a dose-dependent manner (**Figure 5**), with nearly identical activity to M-4H2 IgG. Similarly, western blot analysis also indicated specific interaction between the plant-made chimeric IgG and IgM mAbs and both recombinant CTS1 and MCF (**Figure 6**), further validating the binding activity of the P-4H2 IgG and P-4H2 IgM to the antigen of interest.

**Figure 5:**
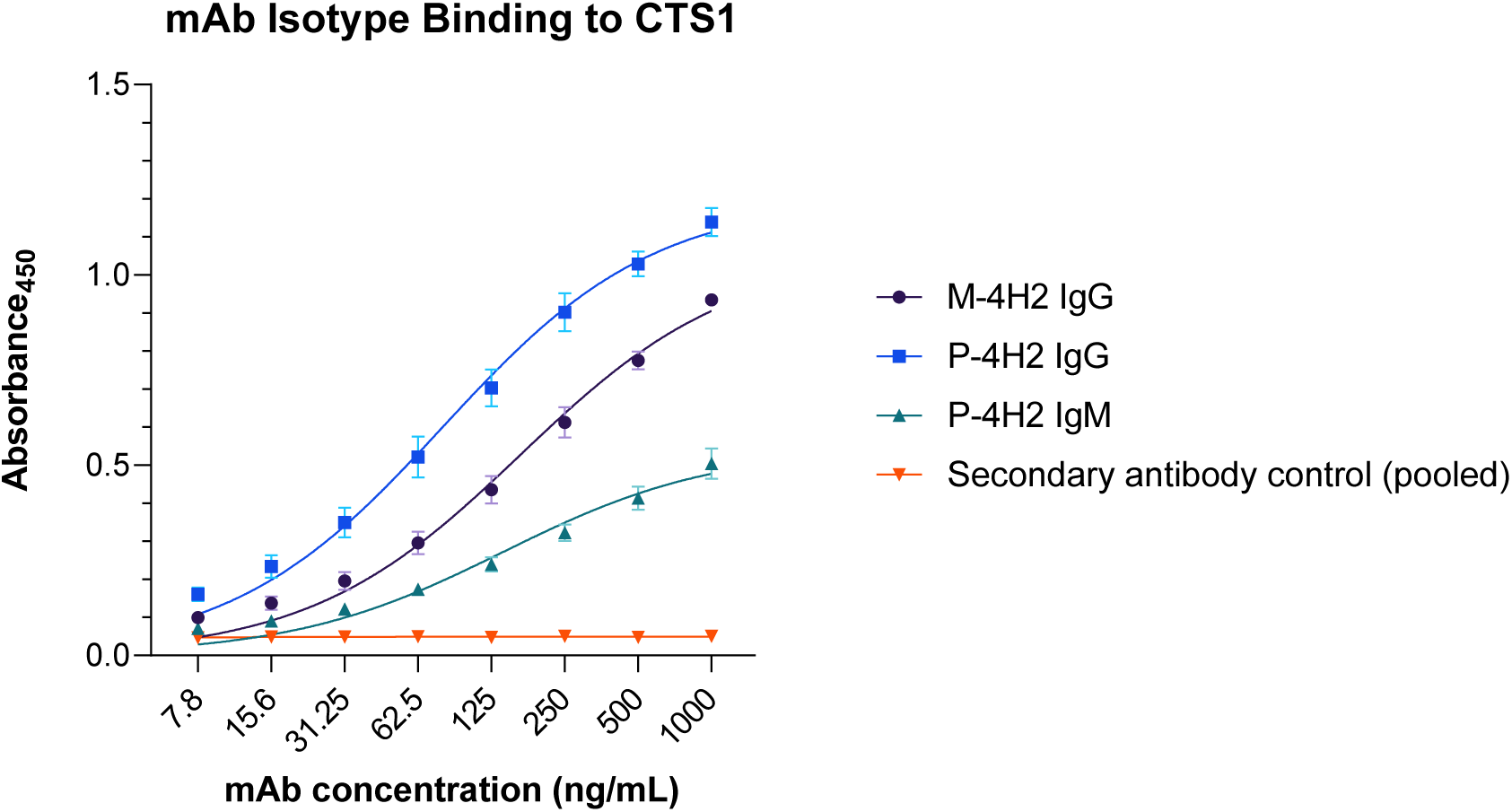
Plant-Made 4H2 Isotypes Specifically Bind to Target Antigen. Serial dilutions of P-4H2 IgG and P-4H2 IgM isotypes were incubated with CTS1 immobilized on ELISA plates. An HRP-conjugated anti-human IgG or anti-human IgM secondary antibody was used to detect the specific interaction between the plant-made IgG or IgM isotype (P-4H2 IgG, P-4H2 IgM) with CTS1, respectively. The parental mouse IgG (M-4H2 IgG) was used as a positive control, which was detected with an anti-mouse IgG secondary antibody conjugated to HRP. Mean ± SEM is plotted from three independent experiments.

**Figure 6.**
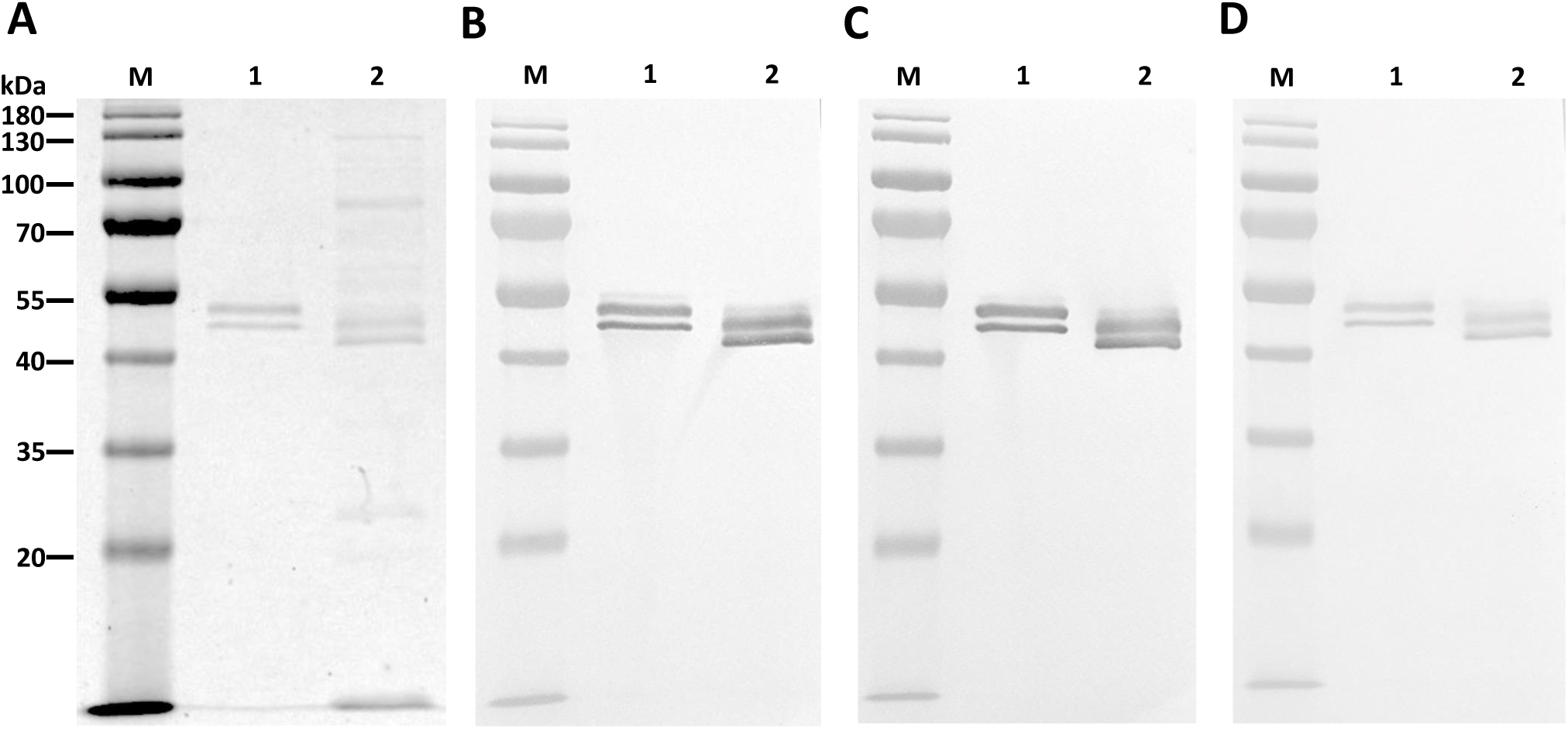

## Discussion

In this study, we utilized plants as an alternative recombinant protein expression platform to express two isotypes of a hybridoma-produced mAb against a fungal antigen, CTS1, from *Coccidioides* spp. To our knowledge, this is only the second report of generating an IgM molecule in plants, but a first for our intentional monomeric IgM design (pairing of two heavy chains), as opposed to a pentameric design^17,32^. Both the humanized P-4H2 IgG and P-4H2 IgM were efficiently expressed within a week of transgene delivery, with levels of 39.95 ± 2.5 and 33.3 ± 1.19 µg/g FLW, respectively. These expression levels fall within the range of previously reported plant-made mAbs, but recombinant IgG levels vary widely, with observations ranging from 4 – 1500 µg/g FLW^33,34^, while a previous expression of a pentameric IgM reached levels of approximately 84 µg/g FLW^17^. The moderate expression levels of the IgG and IgM observed in this study may be attributed to several factors, including the use of non-codon-optimized plant protein expression, specifically in the variable regions of the mAbs, and the use of different expression vectors than those reported in the literature. Increasing expression levels of these mAbs is possible by addressing these points and efforts to increase protein expression levels are currently ongoing.

The sandwich ELISA used to monitor expression level was designed to only recognize recombinant mAbs that contained both a human kappa light chain and either a human gamma (for IgG) or human mu (for IgM) heavy chain. The proper assembly of both the IgG and IgM monomers was further validated by reducing and non-reducing western blot. Indeed, the IgG kappa chain was detected at the expected size of ∼150 kDa. Furthermore, the kappa and mu chain components of the IgM observed in SDS-PAGE were identified and pairing of two mu heavy chains and two kappa light chains to form a monomeric IgM was shown to occur, indicated by the ∼170 kDa band under non-reducing conditions both on western blot and SDS-PAGE. We expected no more than a monomeric form of the IgM, due to the intentional absence of the joining (J)-chain gene, yet a similar band pattern can be observed when comparing the plant-made monomeric IgM with the pentameric human IgM control, indicating possible formation of other IgM oligomers without the J-chain, which has been observed in other studies^35–39^. The smeared banding pattern of the plant-made IgM can be attributed to the presence of heterologous populations of IgM in the purified product, likely due to proteolysis. Optimization of purification of IgM can be performed in the presence of protease inhibitors to limit potential proteolysis. If proteolysis cannot be avoided, the target IgM population could be further purified by other additional chromatography steps.

Both the chimeric P-4H2 IgG and P-4H2 IgM were purified by simple affinity chromatography methods, specific for their respective heavy constant regions. The IgG reached comparable levels of purity to a mammalian cell-produced IgG subjected to the same purification scheme. In contrast, the homogeneity of P-4H2 IgM was lower than the pentameric IgM control made in mammalian cells.

Indeed, the major smaller-than-expected bands between 37 and 50 kDa appeared as a smear of protein fragments, with a minor band appearing just above the 20 kDa marker. We can conclude that the bands of ∼45 kDa in size are degradation products from human mu chain, due to the use of an anti-human mu chain agarose as the affinity resin in purifying the mAb isotype. It is also reasonable to infer that the band near 20 kDa is a mu chain degradation product, given that the summation of the ∼45 kDa and the ∼20 kDa bands would give an expected size of the non-degraded human mu chain we observe of ∼60-65 kDa. Thus, all the major non-target bands found in the purified, recombinant IgM are accounted for and can be linked back to the constant human mu chain.

For ensuring a consistent and reliable IgM product, several steps can be taken to reduce or eliminate the unwanted bands we observed here. Our monomeric design may just be inherently unstable. Including a J chain gene can be implemented to preferentially produce a pentameric IgM product, as has been performed previously in plants^17^, which would likely increase the stability of the protein, help protein folding, and may make it less prone to proteolytic degradation during protein extraction and purification. We can also adjust the purification method to include a second affinity chromatography step specific for human kappa constant chain, potentially only purifying active species of IgM (those that have both mu and kappa chains). This would provide a consistent product for the intended use as an antigen-specific human dilution control.

Valley Fever is a fungal disease that requires an accurate and timely diagnosis. However, current FDA-approved diagnostic kits lack a dilution control. Currently, clinical laboratories who run serologic tests to detect antibodies against *Coccidioides spp*. are forced to identify, collect, store and re-use previously IgG and IgM-positive sera for use in subsequent test runs. This leads to inconsistency in diagnoses due to inherent variability of storing and re-using human sera. In response to this unmet need for a consistent diagnostic control, we developed a mouse mAb specific to a *Coccidioides* spp. antigen used in the clinical diagnostic kits and engineered it into human IgG and IgM chimeras. We confirmed that the human chimeras retain original antigen recognition by indirect ELISA with recombinant CTS1.

This binding was validated in parallel by Western blotting using recombinant CTS1, alongside the immunogen, MCF. Taken together, these data provide evidence that the plant-made chimeric IgG and IgM specifically recognize the antigen of interest found in the immunogen, in a manner consistent with the parental murine mAb. Furthermore, the humanization allows these chimeric mAb isotypes to be recognized by anti-human IgG and IgM secondary antibodies in an ELISA, like those used in Valley Fever diagnostic kits. Therefore, these human chimeric mAbs may be useful as positive controls to replace positive human sera currently used to comply with federal law. This may lead to more consistent diagnostic results and a sustainable supply. The next phase of research will involve investigating the utility of these mAbs in the commercial kits with clinical samples and gaining regulatory approval.

Overall, our study highlights the opportunity of plant-based, recombinant production of both IgG and IgM isotypes against CTS1, as consistent controls for Valley Fever diagnostic kits. This platform may provide a consistent and reproducible source of low-cost mAbs as diagnostic reagents for other diseases as well.

## Author Contributions

Q.C., D. L. and T.G. conceptualized research; C.J. and F.G. designed experiments, performed experiments, and analyzed data. C.J., F.G. and Q.C. wrote the paper with revision by D.L. All authors have reviewed and agreed to the submitted version of the manuscript.

## Acknowledgements

The authors wish to thank Dr. Herta Steinkellner for the sequence of mu constant region. We also thank Katherine Nguyen, Joshua Lesio and Akhil Mahant for helping with *N. benthamiana* maintenance.

## Figure Legends

**Supplemental Figure 1: Plant-Made 4H2 Isotypes Bind to Target Antigen on Western Blot**.

Recombinant CTS1 and *Coccidioides posadasii* mycelial culture supernatant were separated by SDS-PAGE and either stained with Coomassie Blue (**A**) or transferred to PVDF membrane. Membranes were then probed with the parental M-4H2 IgG (**B**), the humanized, P-4H2 IgG (**C**), or the humanized, P-4H2 IgM (**D**), followed by the appropriate secondary HRP-conjugated antibody and development with a colorimetric substrate. Lane 1, Recombinant CTS1. Lane 2, *Coccidioides posadasii* mycelial culture supernatant. M, molecular weight marker.

